# Structure of human cytoplasmic Pol II complex explains global transcription repression by Gdown1

**DOI:** 10.64898/2025.12.10.693371

**Authors:** Jana Schmitzová, Yumeng Zhan, Srinivasan Rengachari, Frauke Grabbe, Olexandr Dybkov, Henning Urlaub, Michael Lidschreiber, Christian Dienemann, Patrick Cramer

## Abstract

RNA polymerase II (Pol II) is a 12-subunit enzyme crucial for gene transcription in the nucleus. However, its assembly in the cytoplasm, nuclear import, and nuclear function of assembly factors remain poorly understood. Here, we isolated Pol II from the cytoplasmic fraction of human cells (cfPol II) and determined its cryo-EM structure. The structure reveals that Pol II is fully assembled in the cytoplasm before nuclear import. We also found that Gdown1 binds Pol II through three distinct regions, indicating it may stabilize Pol II assembly intermediates. Notably, Gdown1 binding precludes the association of essential transcription factors IIB and IIF, rendering cfPol II inactive in promoter-dependent transcription *in vitro*. Our results provide a basis for Gdown1-dependent global transcription repression and suggest a model for the role of Gdown1 in Pol II assembly, import, and transcription regulation.

## Introduction

RNA polymerase II (Pol II) is a 12-subunit complex that is responsible for transcription of protein-coding genes and various regulatory RNAs, including snRNAs, lncRNAs, miRNAs, and eRNAs^1^ across eukaryotes. Pol II function is essential for cell differentiation, development, and environmental responses^2^. The first high-resolution structure of a eukaryotic Pol II was determined over two decades ago^3^ and since then, a large amount of structural and mechanistic data have accumulated. However, the majority of these studies have focused on understanding how Pol II regulates gene expression in the nucleus^4^. On the other hand, the molecular basis underlying Pol II assembly in the cytoplasm, its nuclear import, and the intranuclear release of assembly and transport factors from Pol II remain less well known.

Studies in yeast and subsequent work in human systems have shed light on some key aspects of Pol II biogenesis and assembly^5–8^. Pol II assembly proceeds via two major intermediates centered on its largest subunits – RPB1 and RPB2. First, the formation of an RPB2-containing subcomplex comprising the Pol II subunits RPB2, RPB9, RPB3, RPB11, RPB10, and RPB12 is assisted by the GTPases GPN1, GPN2, GPN3, RNA polymerase II associated proteins 1 and 2 (RPAP1, RPAP2), and Gdown1^9–13^. Second, the RPB1-containing subassembly comprising RPB1, RPB5, RPB6, and RPB8 is assembled and stabilized by heat- shock protein 90 (HSP90) and the R2TP complex^14^. These subassemblies then consolidate with the RPB4/7 dimer to constitute the complete 12-subunit Pol II complex.

We still lack an understanding of the exact stoichiometry of Pol II subunits during nuclear import. However, depletion of individual Pol II subunits results in cytoplasmic accumulation of RPB1, suggesting that the complete assembly of Pol II precedes its nuclear import^7,14^. SLC7A6OS (yeast Iwr1^15^), which likely binds a fully assembled Pol II, facilitates nuclear import of Pol II. Assembly factors that are still associated with Pol II are imported together with Pol II^5,11,12,16,17^.

Among the factors needed for Pol II assembly Gdown1, GPN1, GPN3, and RPAP2 are the most characterized. Gdown1 is a metazoan-specific protein that resides predominantly in the cytoplasm, either in its free form, as part of an RPB2-containing Pol II subassembly, or bound to fully assembled Pol II^7^. Gdown1 is imported to the nucleus together with Pol II^18^. In addition, Gdown1 has been shown to interfere with Pol II transcription in the nucleus at the level of initiation^19–23^, pausing^20^, elongation^20,24^ as well as termination^25^. Additionally, Gdown1 accumulates in the nucleus at the onset of mitosis, where it was proposed to act as a global suppressor of Pol II transcription^18^. GPN1 and GPN3 form a complex that can interact with the CTD of RPB1, and with RPB4/RPB7^11^. The GTPase activity of GPN1 is required for Pol II nuclear localization and both GPN1 and GPN3 are essential for cell viability, underscoring their critical role in Pol II assembly^3,11^. RPAP2 (yeast Rtr1^26^) shuttles between the cytoplasm and nucleus in a GPN1-dependent manner^17^. RPAP2 controls gene expression by modulating Pol II transcriptional activity and regulating nuclear Pol II levels^17^.

To date, structural insights into Pol II regulation by assembly and import factors is limited to high-resolution structures of Pol II-RPAP2 complexes^27,28^, low-resolution models of Gdown1 bound to Pol II^29,30^ in humans and a yeast Pol II-Iwr1 complex^15^. Structures of Pol II in complex with more than one assembly factor are lacking, which limits deeper mechanistic understanding of these factors in Pol II assembly, import and their associated functions after import to the nucleus.

To address this, we have developed a system for the isolation of Pol II from the cytoplasmic fraction (cfPol II) of HEK293 cells expressing N-terminally tagged RPB1. Using biochemical and proteomic characterization of cfPol II, we confirm the co-purification and binding of the well-known Pol II assembly and import factors including Gdown1, GPN1, GPN3, RPAP2 and. Single-particle cryo-EM analysis of cfPol II reveals the first high-resolution structure of human Pol II bound by two assembly factors, namely RPAP2 and Gdown1. The structure shows how Gdown1 binds to the Pol II surface in a multi-valent manner. Comparative structural analysis explains the incompatibility between Gdown1 and transcription initiation factors. Further, we show that cfPol II is inactive in de novo transcription initiation *in vitro*. These findings help to explain how Gdown1 can globally repress transcription and suggest a model for the cellular role of Gdown1.

## Results

### Isolation and cryo-EM studies of human cfPol II

To investigate cfPol II, we purified Pol II complexes from the cytoplasmic fraction of HEK293 cells expressing Pol II that is tagged at the RBP1 N-terminus^31^ (Methods, **Extended Data Fig. 1a**). Biochemical analysis of cfPol II showed that it was bound by several additional proteins (**Fig. 1a**) and using mass-spectrometry, we identified these as GPN1, GPN3, RPAP2 and Gdown1 (**Methods, Extended Data Table 1**). The composition of our cfPol II is consistent with the current model for Pol II assembly in the cytoplasm and subsequent nuclear import^5,7^. Cryo-EM analysis of this sample resulted in a reconstruction with clear density for only Pol II and RPAP2 (data not shown). To improve the homogeneity and stoichiometry of the cfPol II for cryo-EM studies, we added recombinantly purified Gdown1 and RPAP2-GPN1-GPN3 complex and purified the complex by size exclusion chromatography (**Extended Data Fig 1b-d**). This complex was mildly cross-linked to improve stability and then subjected to single-particle cryo-EM (**Extended Data Fig. 1e**, Methods). Analysis of the cryo-EM data (**Extended Data Fig. 2**, **Extended Data Fig. 3**) yielded a cfPol II reconstruction at an overall resolution of 2.9 Å (**Fig. 1b**, **Extended Data Table 2**). Most regions of Pol II are well resolved except for the clamp domain, which is largely absent (**Extended Data Fig. 4a**). This phenomenon is previously observed before for free Pol II^32^ and the Pol II-RPAP2 complex^27,28^. Consistent with the composition of cfPol II (**Fig. 1a**), density for the conserved N-terminus of RPAP2 bound to RPB5 and the RPB1 jaw domain is well resolved (**Extended Data Fig. 4**). RPAP2 positioning in our structure is largely identical to previously determined Pol II-RPAP2 complex structures, with minor deviations^27,28^ (**Extended Data Fig. 4b,c**). We did not identify any corresponding density for GPN1 and GPN3, although biochemical analysis confirmed they are present in the cfPol II sample (**Fig. 1a**) and are known to interact with the Pol II CTD and stalk^11^. While chemical crosslinking coupled with masspectrometry (XL-MS) analysis confirmed the association of GPN1 and GPN3 with reconstituted cfPol II (**Fig. 1c**, **Supplementary Table 1**), the absence of clear cryo-EM density for them suggests a weaker association or structural flexibility relative to Pol II. After unambiguously fitting Pol II and RPAP2, unassigned cryo-EM density was present at the Pol II lobe and protrusion domains (**Extended Data Fig. 5a**).

**Fig. 1:**
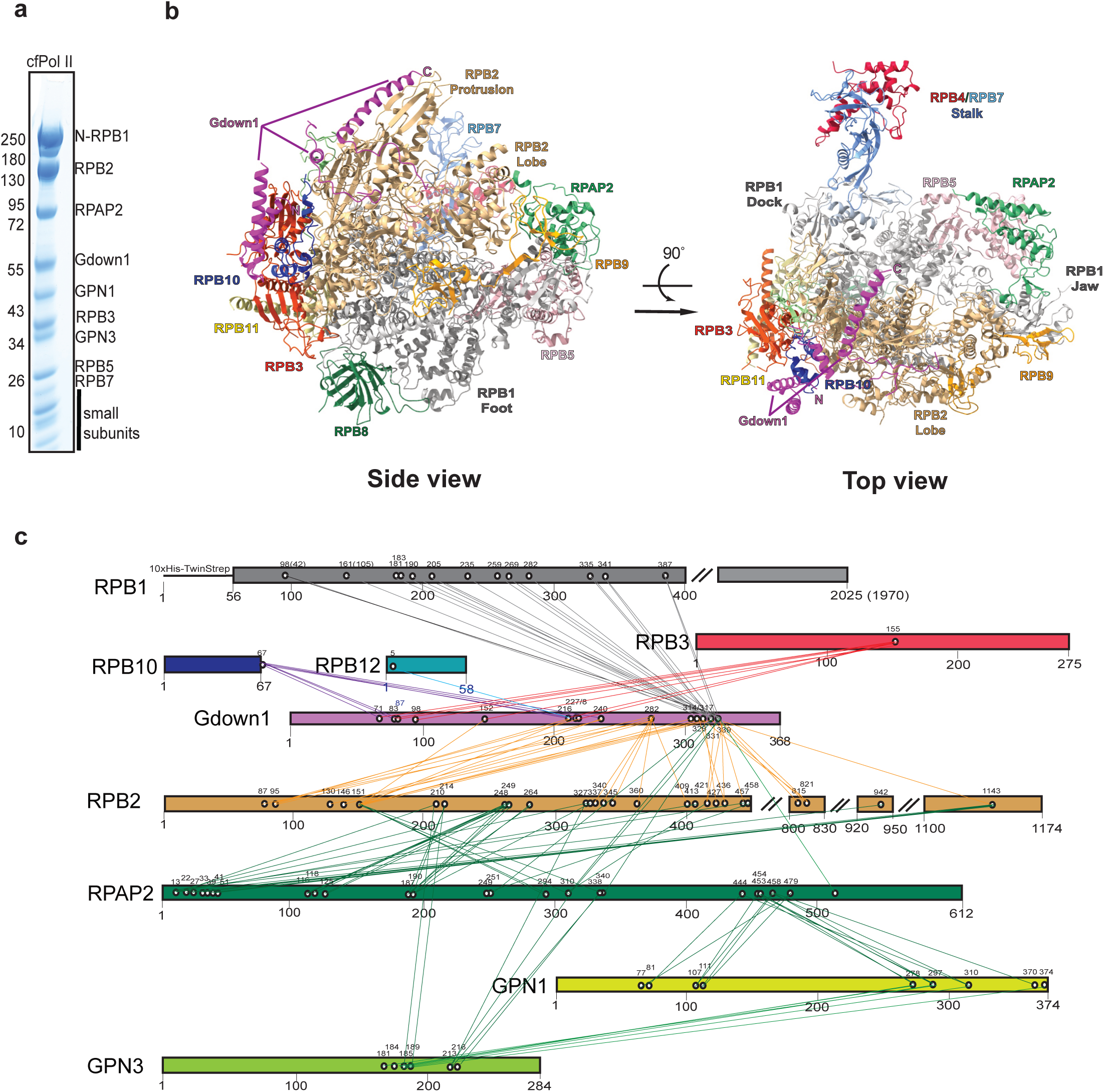
Structure of the human Pol II–RPAP2–Gdown1 complex. **(a)** SDS-PAGE analysis of size-exclusion purified human Pol II supplemented with recombinant Gdown1, RPAP2, GPN1, and GPN3. **(b)** Ribbon representations of the complex. *Left:* Side view of human Pol II highlighting the Gdown1 interaction platform (magenta). *Right:* Top view of human Pol II detailing the RPAP2–Pol II interface. **(c)** Intermolecular crosslinking network of the complex. Diagram shows cross-links between: Gdown1 and Pol II subunits (RPB1, RPB2, RPB3, RPB10, RPB12); RPAP2 and RPB2; and RPAP2 and GPN1/GPN3. Color coding corresponds to panel (b) and is consistent with Cramer *et al*. (2001)^3^.

Previous low-resolution cryo-EM structures for Gdown1 bound to Pol II have localized Gdown1 to the Pol II lobe and protrusion regions^30^. We therefore carefully sorted and locally refined our cryo-EM data to resolve this density to a local resolution range of 2.9-4.2 Å (**Extended Data Fig. 2**). Using this improved cryo-EM map and an Alphafold 3^33^ predicted model of a Pol II-RPAP2-Gdown1 complex, we unambiguously assigned and built Gdown1 residues 19-73, 224-255 and 290-334 into the cfPol II structure. In summary, our purification strategy enabled the preparation of a stochiometric cfPol II complex comprising RPAP2, Gdown1, GPN1 and GPN3 bound to Pol II. Cryo-EM analysis of this sample led to a high-resolution structural model of the Pol II-RPAP2-Gdown1 complex.

### Structure of Gdown1 bound to Pol II

Our high-resolution cryo-EM structure of the Pol II–RPAP2–Gdown1 complex provides the first atomic model of Gdown1. We identified three distinct regions of Gdown1 that bind to Pol II (**Fig. 2a,b**). First, the N-terminal part of Gdown1 (residues 19-73) binds to Pol II subunit RPB10, which we refer to as the RPB10-binding region (RBR, **Fig. 2a,b**). The RBR comprises three α-helices (α1–α3) and includes the conserved LPDKG motif^25^. It binds primarily to RPB10 helices α1 and α2 via hydrophobic interactions (**Fig. 2c**), with additional polar and electrostatic contacts between Gdown1 α1 and RPB3 (**Fig. 2d**). Notably, residues R30 and L44 contribute to the Pol II binding interface, consistent with previous biochemical studies^25,30^. Second, the central part of Gdown1 resolved in our structure (residues 224-255) binds to the Pol II protrusion and external 1 and 2 domains. We refer to this region as external-binding region (EBR, **Fig. 2a,b**). The EBR contains a single α-helix (α4) flanked by unstructured segments. Gdown1 α4 forms hydrophobic contacts with RPB2 helix α3 and RPB12, while the surrounding flexible regions form a mix of hydrophobic, polar, and charged interactions with the protrusion, external and lobe domains of RPB2 (**Fig. 2e**). Third, the C-terminal part of Gdown1 (residues 290-344) binds to the Pol II protrusion domain and we name it protrusion binding domain (PBR, **Fig. 2a,b**). The α5 (residues 305–344) of the PBR anchors Gdown1 to the tip of the protrusion while the less structured stretches bind to the protrusion base (**Fig. 2f**). Residue L303 and L304, previously shown to be critical for transcription inhibition^30^, participates in a hydrophobic interaction network at the protrusion base.

**Fig. 2:**
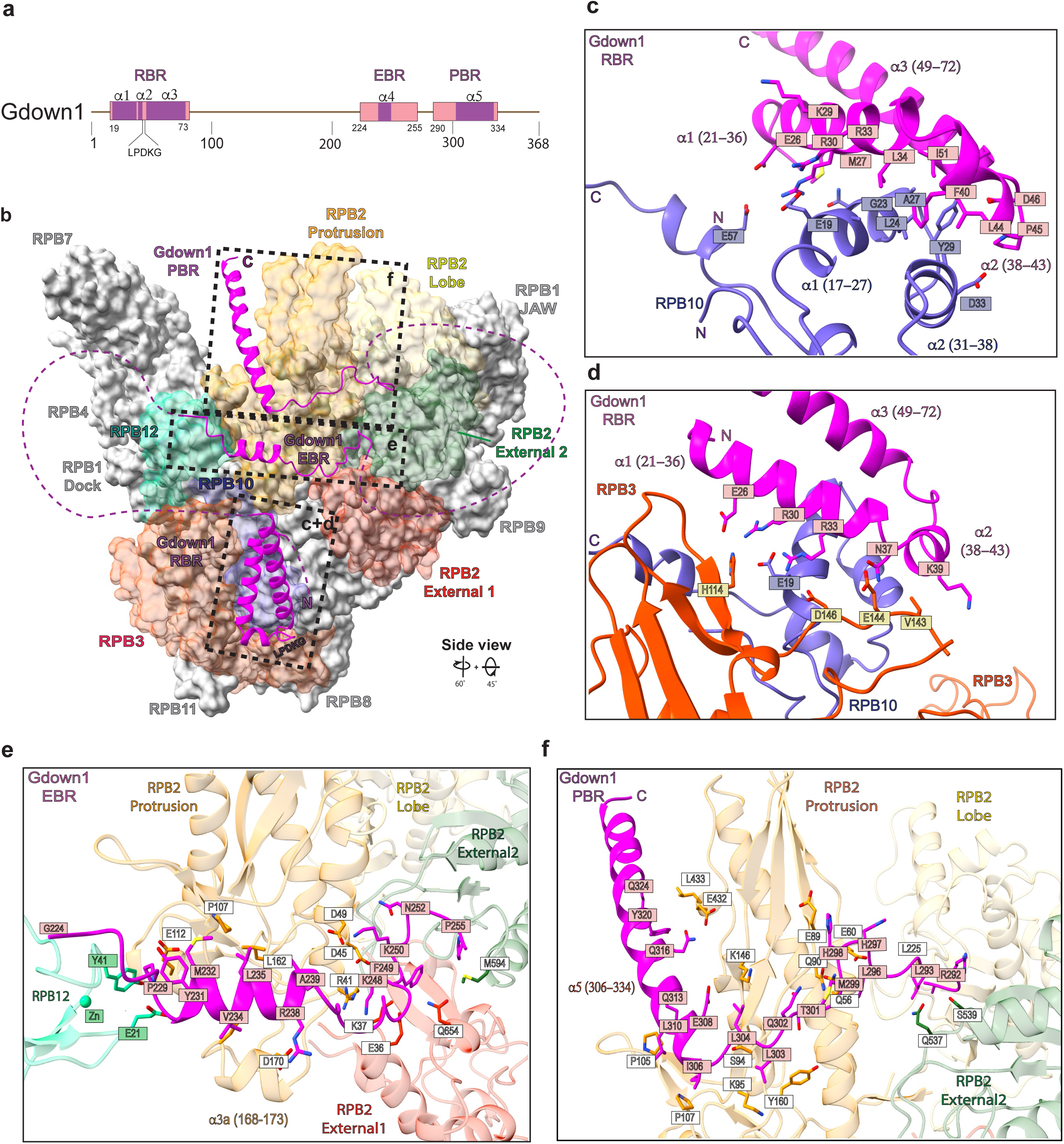
Gdown1 binding interfaces with Pol II subunits. **(a)** Domain architecture of human Gdown1. Modeled regions are shown in magenta: N- terminus (residues 19–73), named as RPB10 Binding Region (RBR); middle region (residues 224–255), named as External Binding Region (EBR); C-terminus (residues 290–334), named as Protrusion Binding Region (PBR). The magenta line indicates disordered regions unresolved in cryo-EM Map A. **(b)** Surface representation of human cfPol II highlighting Gdown1-binding sites. The ribbon model of human Pol II (initial side view orientation) was rotated 60° around the y-axis, followed by a 45° rotation around the x-axis. Domains of RPB2, proteins RPB10, RPB3, and RPB12 follow consistent coloring throughout as in Cramer *et al*, 2001^3^. Gdown1 is shown as a magenta ribbon with unstructured regions marked by magenta dashed lines. **(c)** Interaction interface between Gdown1 RBR (residues 19–73) and RPB10. **(d)** Interaction interface between Gdown1 RBR (residues 19–73) and RPB3. **(e)** Interaction interface between Gdown1 EBR (residues 224–255) and RPB2 (protrusion, external 1 and 2 domains). **(f)** Interaction interface between Gdown1 PBR (residues 290–334) and RPB2 (protrusion domain, external domain 2). Color coding matches Fig. 2b.

The positioning of the RBR, EBR and PBR in our Pol II -RPAP2-Gdown1 structure presents significant deviations from the previous work ^30^ (**Extended Data Fig. 5b,c**). In particular, the RBR comprises residues 19-73, whereas the N-terminal transcription inhibition region (N-TIR) was proposed to range from 1-65. Next, the residue range of the EBR (224- 255) differs from the Pol II binding region I (216-299) that was proposed to mediate Pol II binding. Finally, the RBR (residues 290-334) comprises the previously identified C-terminal transcription inhibition region (C-TIR) and Pol II binding region II, however, our structure shows they rather belong to a single region. Altogether, our high resolution cryo-EM structure of Gdown1 allowed us to build the first atomic model for Gdown1 bound to the Pol II surface. We resolved three distinct structural regions of Gdown1 that bind to Pol II in a multivalent manner.

### Gdown1 binding to Pol II overlaps with transcription initiation factors

Multiple studies have shown Gdown1 to compete with transcription initiation^19,20,30^ factors to bind Pol II, thereby interfering with transcription in the nucleus^21,22^. To explore the structural basis underlining these phenomena, we compared our high-resolution model of Gdown1 bound to Pol II to the structure of the pre-intiation complex (PIC) ^34^. The comparison identiefies three main clashes between Gdown1 and general transcription factors (GTFs). First, the Gdown1 PBR overlaps with the TFIIB cyclin domains (**Fig. 3a**). Second, the α5-helix of the PBR extends into the trajectory of promoter DNA (**Fig. 3a**). Third, the path of the PBR on the Pol II surface largely overlaps with the linker between the dimerization domain and the C-terminal WH domain of TFIIF-β (**Fig. 3b**). Moreover, although the region between the PBR and EBR is disordered, it might sterically interfere with binding of TFIIF to the Pol II lobe (**Fig. 3b**). These clashes are consistent with studies showing that Gdown1 competes with TFIIB and TFIIF in the PIC and inhibits transcription initiation^19,20,22,30^. Altogether, our structure of the Pol II-RPAP2-Gdown1 complex shows how Gdown1 binding to Pol II may preclude transcription initiation factors from binding Pol II and thus explains how Gdown1 can inhibit PIC assembly and transcription initiation.

**Fig. 3:**
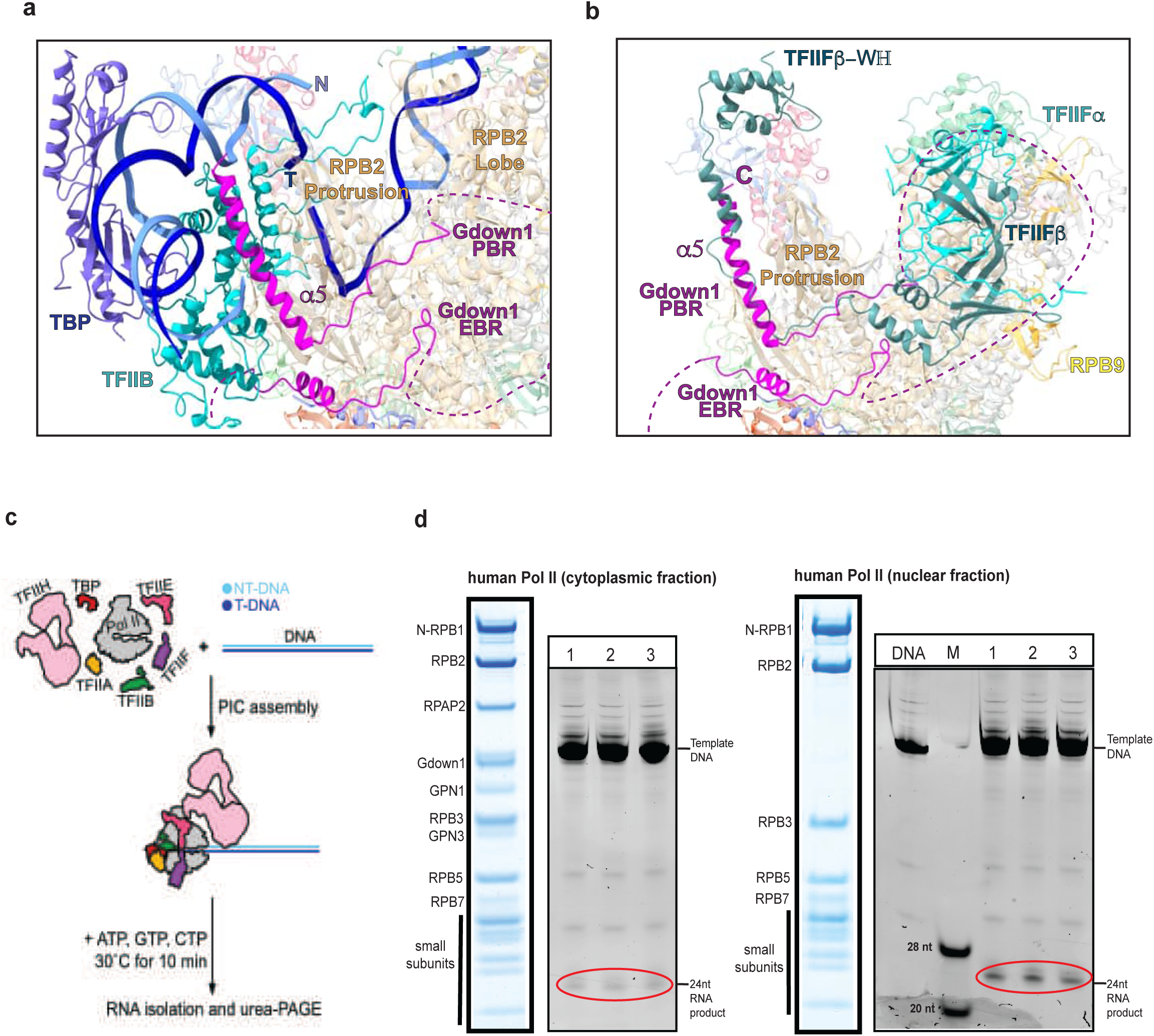
Gdown1 binding to Pol II sterically clashes with transcription initiation machinery. **(a)** Structural clash analysis. Gdown1 PBR (magenta ribbon) overlaps with transcription initiation factors TBP (blue) and TFIIB (cyan), and promoter DNA. A Pol II pre-initiation complex (PIC) model (PDB: 8S52^34^) was superimposed on our Pol II-RPAP2-Gdown1 structure. Pol II is shown at 80% transparency. Color scheme matches Fig. 1. **(b)** Reduced transcriptional activity of cytoplasmic Pol II. *Top:* Schematic of in vitro transcription initiation assay. *Bottom:* Cytoplasmic Pol II shows diminished initiation activity compared to nuclear Pol II. NT-DNA: non-template strand; T-DNA: template strand. **(c)** Schematic of de novo transcription initiation assay performed in vitro. **(d)** Biochemical validation. *Left:* SDS-PAGE of nuclear-tagged vs. cytoplasmic Pol II preparations. *Right:* Urea-PAGE of transcription products. M: RNA marker (28 nt and 20 nt bands). Red circle indicates initiation product. Experiments performed in triplicate (n=3). Nuclear Pol II was isolated as previously described³¹; cytoplasmic Pol II was prepared as in Extended Data Fig. 1a and Methods.

### Purified cfPol II containing Gdown1 is transcriptionally inactive

The structural incompatibility between Gdown1 and parts of the Pol II PIC indicate that cfPol II may be unable to initiate transcription. To test whether cfPol II is able to initiate transcription *in vitro*, we performed *in vitro* transcription initiation assays using promoter DNA and purified GTFs (Methods, **Fig. 3c**). To monitor transcription initiation while reducing the influence of altered transcription elongation, transcription reactions were performed using promoter DNA containing a 24 bp U-less cassette would have less effect.

When purified cfPol II was used in the transcription initiation reaction, only a weak signal for the expected 24 nt RNA product could be detected (**Fig. 3d**). In contrast, Pol II that was purified from the nuclear fraction of the same cell line initiates transcription *de novo* and RNA synthesis was observed (**Fig. 3d**). This finding is consistent with Gdown1 competing with GTFs and promoter DNA, thereby inhibiting transcription. Thus, cfPol II that is bound by Gdown1 is unable to initiate transcription.

## Discussion

In conclusion, our structure of cfPol II provides the first atomic model for Gdown1 bound to Pol II, and shows three distinct structural regions of Gdown1 that bind to RPB10 as well as the external and protrusion domains of Pol II. Gdown1 binding to Pol II sterically clashes with transcription initiation factors TFIIB and TFIIF as well as promoter DNA, and cfPol II is unable to initiate transcription *in vitro*. Together with previous studies, our findings establish the basis for Gdown1-dependent transcription repression and suggest a model for how Gdown1 may facilitate Pol II assembly and import, and may rapidly be exported upon dissociation from Pol II.

During Pol II assembly, Gdown1 binds the subcomplex comprising RPB2, RPB3, RPB10, RPB11 and RPB12^14^, and binding of Gdown1 to cfPol II bridges between Pol II subunits RPB2, RPB10 and RPB3 (**Fig. 2b**). Based on similarity to the assembly of bacterial RNA polymerase, RPB2 and RPB3 have been hypothesized to be part of separate subcomplexes that then associate to form the RPB2, RPB9, RPB3, RPB10, RPB11 and RPB12 subassembly at an early step of Pol II assembly^7^. We therefore propose that Gdown1 may stabilize formation of this subcomplex and thereby functions during cytoplasmic Pol II assembly.

Gdown1 binds to Pol II before nuclear import, as evident from our isolation of cfPol II. The Gdown1 C-terminus contains a cytoplasmic anchoring signal (CAS), that was shown to facilitate localization to the nuclear pore by interacting with RAE1 and NUP50 at the cytoplasmic and nuclear side of the pore, respectively^2^ . This indicates that Gdown1, when bound to cfPol II, can facilitate nuclear import of Pol II.

Gdown1 is imported to the nucleus alongside with Pol II^7^ and nuclear accumulation of Gdown1 was shown to correlate with low transcriptional activity^30^ and cause global repression of transcription^2^. Our structural and functional data explain how Gdown1 acts as a repressor of transcription by interfering with PIC assembly. Moreover, the Gdown1 binding site on Pol II also overlaps with the binding sites of elongation, pausing and termination factors PAF1, LEO1, RTF1, NELF-A, ELL2 and TTF2 (**Extended Data Fig. 6**), further suggesting that Gdown1 can also interfere with the nuclear function of these factors.

Altogether our structural and biochemical data substantiates the role of Gdown1 as a global repressor of transcription in the nucleus.

To render nuclear Pol II transcriptionally active, Gdown1 needs to be released and exported to the cytoplasm where the majority of Gdown1 localizes^2,18^. Recent studies showed that phosphorylation of Gdown1 at S270^25^, likely by the Mediator kinase CDK8, relieves the inhibition of transcription initiation^19,23,35^. In our structure S270 resides in the unresolved part between the EBR and PBR, which precludes from understanding how its phosphorylation regulates Pol II association. Gdown1 comprises two nuclear export sequences (NES)^18^ that facilitate rapid nuclear export of free Gdown1. One of the two NES resides at the C-terminal end (residues 332-340) of the helix α5 of PBR2 and interacts with the Pol II protrusion (**Fig. 2**). This NES thus may only become available upon Gdown1 release from Pol II, indicating tight regulation between Gdown1 release from Pol II and Gdown1 nuclear export.

Altogether, our structural and functional data propose a model by which Gdown1 may facilitate Pol II assembly and import, acts as a global transcription repressor in the nucleus, and may rapidly be exported upon dissociation from Pol II. Our work establishes the structural framework to study Gdown1 function and paves the way for future studies investigating the molecular mechanisms by which Gdown1 controls Pol II assembly and nuclear import, and facilitates transcription repression in the nucleus.

## Methods

### Molecular cloning

The cDNAs encoding full-length human assembly and transport factors GPN1 and GPN3 were amplified from a human cDNA library using following primers gpn1_fwd: 5′- TACTTCCAATCCAATGCAATGGCGGCGTCCGCAG-3′, gpn1_rev: 5′- TTATCCACTTCCAATGTTATTATTTATTGTTTCTCTTCCAGTATTGTGCCATCG-3′, gpn3_fwd: 5′-TACTTCCAATCCAATGCAATGCCTCGGTATGCGCAGCT-3′, gpn3_rev: 5′-TTATCCACTTCCAATGTTATTATTTATTGTTTCTCTTCCAGTATTGTGCCATCG-3′. Amplified GPN1 and GPN3 sequences were cloned into the p438C vector that encodes proteins with an N-terminal His₆-MBP tag.

### Purification of Pol II from the cytoplasm

All protein purification steps were performed at 4°C unless otherwise specified. Pol II was purified from the cytoplasmic fraction of HEK293 cells stably expressing N-terminally His_10_-Twin-Strep-tagged human Pol II subunit RPB1, generated via CRISPR/Cas9 genome editing^31^. Cells were grown in a 15 L fermenter (Applikon BioBundle, 15 L capacity, ez2- control) to a density of 3–6 × 10⁶ cells/mL with viability between 86% and 93%.

The cytoplasmic fraction was prepared according to the method of Dignam et al.^36^. After harvesting, cells were washed twice with 1× PBS buffer and pelleted by centrifugation at 836 ×g for 10 min. The cell pellet was resuspended in hypotonic 1× MC buffer (10 mM HEPES-KOH, pH 7.6; 10 mM KOAc; 0.5 mM Mg(OAc)₂; 0.5 mM DTT), supplemented with EDTA-free protease inhibitors (Roche; 2 tablets per 50 mL buffer), and incubated on ice for 5 min. Swollen cells were lysed using a 100 mL Dounce homogenizer (Kimble Chase, Sigma-Aldrich) with 18 strokes, followed by centrifugation at 18,000 × g for 5 min. The supernatant (cytosolic extract) was flash-frozen and stored at –80 °C.

Prior to purification, the thawed sample was cleared by centrifugation at 26,074 × g for 30 min (rotor Fiberlite F14 6x250y, Thermo Fisher Scientific). To remove nucleic acid contaminants, the supernatant was treated with 5% polyethylenimine (PEI, pH 7.9) to a final concentration of 0.04%. After 10 min of stirring at 4 °C, the resulting white precipitate was pelleted by centrifugation at 18,668 × g for 20 min. The pellet was washed with A0 buffer (50 mM Tris-HCl, pH 7.9; 5 mM MgCl₂; 2 mM DTT; 10% glycerol) using 5× Dounce homogenization followed by centrifugation at 18,668 × g for 20 min. The washed pellet was resuspended in buffer A100 (50 mM Tris-HCl, pH 7.9; 1.5 mM MgCl₂; 2 mM DTT; 10% glycerol; 100 mM ammonium sulfate) by stirring for 60 min at 4°C.

Ammonium sulfate precipitation was performed by adding finely ground ammonium sulfate to achieve 50% saturation. The mixture was stirred at 4 °C for 1 hour and cleared by centrifugation at 25,000 rpm (rotor A27 6x50, Thermo Fisher Scientific) for 60 min. The supernatant was filtered through a 45 μm Filtropur S filter (Sarstedt) and loaded onto a 20 mL self-packed Strep-TactinXT 4Flow column (IBA Life Sciences), which had been equilibrated in buffer A100. The column was washed with 10 column volumes of buffer A100 and eluted with 50 mM biotin (IBA Life Sciences) in buffer A100. All buffers were supplemented with protease inhibitors: 1 mM PMSF, 1 mM benzamidine, 60 μM leupeptin, and 200 μM pepstatin.

The eluted fractions were analyzed by SDS-PAGE using a 4–12% Bis-Tris NuPAGE gel in 1× MES running buffer (Invitrogen). Fractions containing cfPol II associated with assembly factors were pooled and concentrated using Amicon Ultra centrifugal filters (100 kDa molecular weight cutoff, MilliporeSigma). From approximately 32× 10^9^ cells, about 150 μg of cfPol II were purified. Samples for cryo-EM analysis, with or without spike-in of recombinantly expressed and isolated assembly factors, were further purified by size-exclusion chromatography on a Superose 6 Increase column (3.2 × 300 mm, GE Healthcare). Fractions corresponding to monomeric Pol II complexed with assembly factors were pooled, concentrated, and used for further experiments.

### Purification of the RPAP2, GPN1 and GPN3

To facilitate expression of the individual N-terminal His₆-MBP-tagged proteins (RPAP2^27^, GPN1, and GPN3), bacmids were generated using DH10αEMBacY cells. SF9 cells (Thermo Fisher Scientific) were transfected with the corresponding bacmids to produce V0 virus, which was then used to generate V1 virus in SF9 cells. Large-scale co-expression of RPAP2-GPN1-GPN3 was carried out in Hi5 cells (Expression Systems) via co-transfection with V1 viruses encoding each protein subunit. The insect cells expressing the recombinant proteins were harvested at 85–90% viability to minimize degradation.

Frozen cell pellets were thawed at room temperature, resuspended in the lysis buffer A250 (50 mM Tris-HCl, pH7.9; 250 mM ammonium sulfate; 2.5 mM MgCl_2_; 10% glycerol; 0.1 mM EDTA, pH8.0; 2 mM DTT; and protease inhibitors: 1 mM PMSF, 1 mM benzamidine, 60 μM leupeptin, and 200 μM pepstatin), and lysed by sonication.

After centrifugation at 25,000 rpm, for 60 min (rotor A27 6x50, Thermo Fisher Scientific), the supernatant was filtered sequentially through 0.8 μm and 0.45 μm Filtropur S filter (Sarstedt) to remove aggregates. The clarified lysate was then incubated with 10 ml amylose resin (NEB) in batch, pre-equilibrated with lysis buffer A250. The complex was eluted by on-resin cleavage with TEV protease overnight at 4°C. The resulting assembled heterotrimeric complex was further purified on a Sepharose 200 10/300 column (Cytiva), equilibrated in buffer A150 (HEPES-NaOH pH7.5, 150 mM NaCl, 2 mM DTT, 5% glycerol).

### Purification of full-length human Gdown1

Human N-terminal His₆ tagged Gdown1 was expressed from plasmid pOPINB_plus_GDOWN1_FL^32^ in *E. coli* BL21-RIL cells by induction at OD_600_=1.2 with 1 mM IPTG for 18 h at 18°C. Frozen cell pellets were thawed at room temperature, resuspended in the lysis buffer A400 (50 mM HEPES-NaOH, pH7.5; 400 mM NaCl; 30 mM imidazol; 10% glycerol; 2 mM BME; and protease inhibitors: 1 mM PMSF, 1 mM benzamidine, 60 μM leupeptin, and 200 μM pepstatin), and lysed by sonication. The lysate was cleared by centrifugation at 25,000 rpm (rotor A27 6x50, Thermo Fisher Scientific) for 60 min and filtered with 0.45 μm Filtropur S filter (Sarstedt) to remove aggregates. The clarified supernatant was loaded onto a 5 ml HisTrap HP column (Cytiva), pre-equilibrated with lysis buffer A400. His_6_-Gdown1 was eluted with linear imidazole gradient from 30 mM to 400 mM in buffer A200 (25 mM HEPES-NaOH, pH7.5; 200 mM NaCl; 10% glycerol; 2 mM BME). Fractions containing His_6_-Gdown1 were pooled, and the His_6_ tag was removed by incubation with TEV protease overnight at 4°C during dialysis against buffer A100 (25 mM HEPES-NaOH, pH7.5; 100 mM NaCl; 10% glycerol; 2 mM BME). The tag-free Gdown1 was further purified by reverse affinity chromatography: the dialysed sample was incubated with Ni-NTA resin (Qiagen) in batch to remove the cleaved His_6_-tag and residual His_6_-tagged proteins. The flow-through, enriched in tag-free Gdown1, was then applied to a 5 ml Q -Sepharose HP column (Cytiva), equilibrated in buffer A100. Proteins were eluted using a linear NaCl gradient from 0.1 to1 M. Fractions containing pure, contaminant-free Gdown1 were pooled, concentrated and flash-frozen for storage at -80°C.

### Cryo-EM sample preparation

Approximately 150 pmol of cfPol II were incubated with a 2-fold molar excess of recombinantly expressed, tag-free assembly factors (RPAP2, GPN1, GPN3, and Gdown1) for 5 minutes at 4 °C. The resulting complex was purified by size-exclusion chromatography on a Superose 6 Increase column 3.2/300 (GE Healthcare), pre-equilibrated in buffer A150 (20 mM HEPES-NaOH, pH 7.5, 150 mM NaCl, 0.5 mM TCEP, 5% glycerol). Peak fractions enriched in cfPol II containing the spiked-in assembly factors were pooled from four independent runs and concentrated to 1.56 μM. A 100 μL aliquot of the sample was further crosslinked with 0.75 mM BS3 (bis(sulfosuccimidyl)suberate; Thermo Fisher Scientific) and incubated on ice for 30 minutes. The reaction was quenched by adding pH adjusted Tris to 100 mM final concentration, followed by an additional 1-hour incubation at 4 °C. The sample was then buffer-exchanged into A150 (without glycerol) by dialysing for 5 hours at 4°C using a 10,000 MWCO Slide-A-Lyzer MINI Dialysis Unit (Thermo Fisher Scientific). The resulting crosslinked CMPol II complex (Pol II-RPAP2-GPN1-GPN3-Gdown1) was used for cryo-EM grid preparation. A volume of 3.5 μL of the diluted sample (∼600 nM) was applied to one side of freshly glow-discharged R2/2 UltrAuFoil 200 grids (Quantifoil). Grids were blotted for 5 seconds at a blot force of 5, at 4 °C and 98% humidity, and rapidly vitrified by plunging into liquid ethane using a Vitrobot Mark IV (Thermo Fisher Scientific).

### Crosslinking-masspectrometry

75 pmol of the purified cfPol II complexes (without spike-in of the assembly factors) were crosslinked with either 0.7 or 1.4 mM BS3 (Thermo Fisher) for 30 min at 24°C in 50µl of the buffer composed of 20 mM HEPES-NaOH, pH 7.5, 150 mM NaCl, 1.5 mM MgCl2, 2mM DTT. The crosslinking reactions were quenched with 55 mM Tris-HCl, pH 7.5 for 15 min at 24°C. The samples were subsequently denatured with 4M urea, reduced with 5 mM DTT and alkylated with 17 mM iodoacetamide. The samples were then diluted with 50 mM ammonium bicarbonate to reduce urea concentration to 1M and digested with trypsin (Promega, sequencing grade) in a 1:20 enzyme-to-protein ratio (w/w) at 37 °C overnight.

Peptides were reverse-phase extracted using SepPak Vac tC18 1cc/50mg (Waters), eluted with 50% acetonitrile (ACN) / 0.1% trifluoroacetic acid (TFA) and dried in a vacuum concentrator (Eppendorf). Obtained peptides were dissolved in 40 µl of 2% ACN / 20 mM ammonium hydroxide and separated by reverse phase HPLC at basic pH using an xBridge C18 3.5µm 1x150mm column (Waters) at a flow rate of 60 µl/min at 24°C. A gradient of buffers A (20 mM ammonium hydroxide, pH 10) and B (80% ACN / 20 mM ammonium hydroxide, pH 10) was used as mobile phase. Peptides were bound to a column pre-equilibrated with 5% buffer B and eluted over 64 min using a multistep gradient of 5-45%B. Fractions of 60 µl were collected. Peptides eluted between minute 10 and 56 were pooled with a step of 15 min, vacuum dried and dissolved in 5% ACN / 0.1% TFA for a subsequent uHPLC-ESI-MS/MS analysis. The fractions were injected into a Dionex UltiMate 3000 uHPLC system coupled to an Orbitrap Exploris 480 mass spectrometer (both Thermo Fischer) and measured 4 times with a 75 min method. uHPLC was equipped with a C18 PepMAP 100 cartridge (0.3 x 5 mm, 5 μm, Thermo Scientific) and a custom 30 cm C18 main column (75 µm inner diameter packed with ReproSil-Pur 120 C18-AQ beads, 3 µm pore size, Dr. Maisch GmbH). Mobile phase was formed using buffers A (0.1% formic acid) and B (80% ACN / 0.08% formic acid). Peptide were separated by applying a linear multistep gradient of 10-52%B. MS settings were as follows: MS1 resolution, 120000; MS1 scan range, 350-1550 m/z; MS1 normalized AGC target, 300%; MS1 maximum injection time, 25ms; cycle time (Top Speed), 3 s; intensity threshold, 1E4; MS2 resolution, 30000; isolation window, 1.6 Th; normalized collision energy, 28%; MS2 AGC target, 75%; MS2 maximum injection time, 128 ms. Only precursors with a charge state of 3-8 were selected for MS2 using a dynamic exclusion of 20 s.

Protein composition of the samples was determined in a MASCOT search. Using a spectrum counting approach (the number of observed peptide-spectrum matches normalized to the protein size), a restricted database was created to encompass 240 most abundant proteins. This database was used for a protein-protein crosslink search of Thermo raw files with pLink2.3.11 software (https://pfind.ict.ac.cn/se/plink/)^37^. The pLink results were filtered at false discovery rate (FDR) of either 1 or 5% and presented in Supplementary Table1. For model building, a maximum distance of 30 Å between the Cα atoms of the crosslinked lysines was allowed.

### Cryo-EM data collection and processing

Cryo-EM data were collected on a 300 kV Titan Krios transmission electron microscope (Thermo Fisher Scientific) equipped with a K3 direct electron detector (Gatan) and a Quantum LS energy filter. Data were acquired in electron counting mode at a nominal magnification of ×81,000, corresponding to a physical pixel size of 1.05 Å/pixel. A 20 eV energy filter slit was used in EFTEM mode. Automated data collection was performed using SerialEM^38^. Exposure time was 2.316 seconds, with a dose rate of 19.0 e⁻/pixel/s (17.23 e⁻/Å²/s), resulting in a total dose of 39.91 e⁻/Å². The defocus range was set between −0.5 and −2.0 μm. A total of ∼11,000 micrographs were collected.

All movie stacks were processed on-the-fly using Warp^39^ for motion correction, dose weighting, and contrast transfer function (CTF) estimation. Micrographs with poor CTF estimates were excluded from further analysis. Approximately 4.1 million particles were extracted using a box size of 280 pixels (binned 2×, resulting in a pixel size of 2.1 Å/pixel) in RELION 5.0^40,41^.

Particles were subjected to iterative rounds of 2D and 3D classification in CryoSPARC ^42^. Classes showing well-defined features of Pol II and assembly factors, including the Pol Ii stalk, the RPAP2 N-terminal domain, and the C-terminal region of Gdown1, were selected for further processing. These particles (∼2.4 million) were re- extracted in their original, unbinned form (pixel size: 1.05 Å/pixel) and further processed in RELION 5.0. Iterative rounds of CTF refinement, 3D refinement and Bayesian polishing were performed, resulting in a final 3D reconstruction of the Pol II core at a resolution of 2.3 Å.

To improve the resolution for the density corresponding to Gdown1, a soft mask encompassing protrusion, lobe and wall of RPB2, Gdown1 C- and N-termini, RBP10, RPB12 and part of RPB3 was applied to focus 3D classification on 2.4 million particles without image alignment using blush function and with regularization parameter T=4 (**Extended Data Fig. 2**). The 3D refinement of the best class containing 265,000 particles (11%) showing C-terminal and partially N-terminal part of Gdown1 was performed. This map was further used for new soft mask preparation encompassing N-terminal Gdown1, RPB10, RPB12, C-terminal part of RPB3 as well as parts of N-termini, lobe and wall of RPB2, respectively. New focus 3D classification without image alignment using blush function and with regularization parameter T=4 with this soft mask was performed on 2.4 million particles. The best class, containing 472,100 particles was further 3D refined obtaining a reconstruction of the map with 2.6 Å resolution of the core Pol II. Another round of focus 3D classification was performed to get better map for the N-termini of Gdown1. Here, two classes, soft mask on the region of N-termini of Gdown1were used. Further 3D refinement of the best class (188,000 particles) a 2.7Å reconstruction was obtained. To improve the resolution of the C-terminus of Gdown1 a soft mask for the region corresponding to RPB12, of protrusion, lobe and wall of RPB2 was used for focus classification with two classes, blush and T=4. The best class (47, 500 particles) was 3D focus refined resulting in 2.9 Å resolution of the core Pol II (Map A). Resolution ranges for the map regions corresponding to the Gdown1 N-terminus (amino acids 19–73), the central part (amino acids 224–255), and the C-terminus (amino acids 290–334) were 3.3–4.1 Å, 3.1–3.4 Å, and 2.9–4.7 Å, respectively.

The resolution of the N-terminal pat of RPAP2 in Map A was limited to 3.8-5.3 Å. Consequently, we performed additional processing focused on RPAP2, resulting in Map B (Extended Data Fig. 3). We conducted focus 3D classification using a mask encompassing the N-terminal part of RPAP2 and the jaw domain of RPB1 on 2.4 million particles without image alignment applying blush regularization and two classes. The class exhibiting well-defined N-terminal RPAP2 density (584,000 particles) was subjected to global 3D refinement. A second round of 3D classification under identical masking and parameters using three classes yielded a class (214,000 particles) that was refined to Map B. This map achieved a resolution range of 3.1–4.2 Å around the N-terminal RPAP2 and 2.6 Å for the core Pol II.

### Model building

For model building, Pol II from the Pol II dimer model (PDB: 7OZN^43^) was rigid-body docked into Map A in Chimera^44^. The starting models for full length Gdown1 and RPAP2 (residues 1-401) bound to Pol II (RPB1 with residues 1-1546; all other subunits of Pol II are full length) were generated by AlphaFold 3^33^. To model RPAP2, the initial model was docked into Map B, which has better density for RPAP2, and manually rebuilt in Coot^45^and real-space refined with Phenix^46^. The refined RPAP2 model and the starting models of Pol II and Gdown1 were fitted as rigid bodies into Map A. The Pol II-RPAP2- Gdown1 model was then manually adjusted in Coot^45^ against Map A, followed by consecutive rounds of real-space refinement in Phenix^46^ and rebuilding with Coot^45^. The final Pol II-RPAP2-Gdown1 model showed good stereochemistry as validated by Molprobity^47^.

### In vitro transcription initiation assay

General transcription factors and thymus Pol II were purified as previously described^32,48–51^. In vitro transcription initiation assays were performed as previously described^34,52^. In brief, promoter templates were generated by large-scale PCR, followed by ion-exchange chromatography and isopropanol precipitation. For each reaction, 1.1 pmol Pol II, 5.8 pmol TFIIF, 1.2 pmol DNA (or nucleosome), 3.8 pmol TBP, 7.7 pmol TFIIA, 3.8 pmol TFIIB, 1.7 pmol TFIIE and 1.7 pmol TFIIH were used. PIC was assembled in a reaction buffer containing 20 mM HEPES-KOH pH 7.5, 109 mM KCl, 3% glycerol, 6 mM MgCl2, 0.5 mM DTT at 30°C for 60 min. Transcription was initiated with the addition of 0.5 mM ATP, 0.5 mM CTP and 0.5 mM GTP and allowed to proceed for 10 min at 30°C. RNAs were purified by proteinase K digestion and isopropanol precipitation and analysed on an urea-gel (7 M urea, 1x TBE, 20% acrylamide:bis-acrylamide 19:1).

## Data availability

The cryo-EM reconstructions and final model were deposited with the Electron Microscopy Data Base (EMDB) under accession code EMD-XXXX and with the Protein Data Bank (PDB) accession YYYY.

## Acknowledgements

We thank Monika Raabe for performing initial mass spectrometry analysis of purified cfPol II. We are grateful to Ulrich Steurwald for maintaining the cryo-EM infrastructure and to Ute Neef and Petra Rus for maintaining the insect cell facility. We also thank Sara Ahrari, Julio Abril-Garrido, Taras Velychko, James L. Walsche, and Isaac Fianu for productive scientific discussions. P.C. was supported by the Max Planck Society.

## Author contributions

J.S. and P.C. conceived and planned the study; J.S. designed/performed biochemical/cryo- EM experiments (under supervision of C.D.) and interpreted data; J.S. purified assembly/transport factors and cfPol II; Y.Z. designed/performed in vitro transcription initiation experiments and modeling; S.R. supervised initial cryo-EM data processing; F.G. purified GTFs; O.D. and H.U. performed crosslinking mass spectrometry; P.C. acquired funding; J.S. and C.D. wrote the manuscript with input from all authors.

## Extended Data Figure Legends

**Extended Data Fig. 1:**
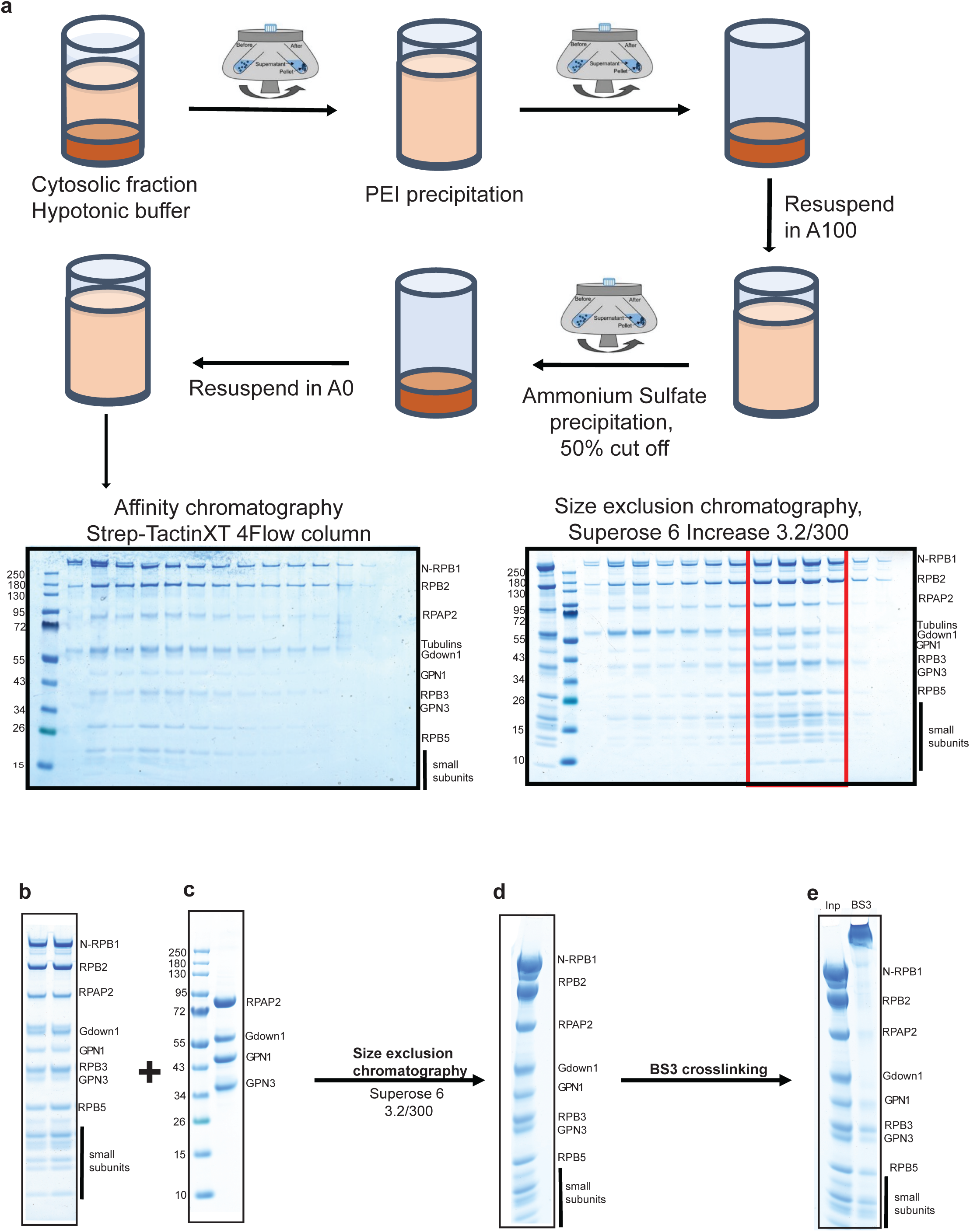
Isolation of human cytoplasmic RNA polymerase II for cryo-EM analysis. **a**, Schematic of cytoplasmic human Pol II purification. Cytosolic fractions from HEK293 cells (clone C1N1 with N-terminal His10-StrepTwin tag on RPB1) were prepared following Dignam et al.^36^. Purified Pol II quality was analysed by SDS-PAGE in MES buffer to resolve all small subunits. Gels were Coomassie blue-stained. After affinity chromatography, Pol II-containing fractions were pooled, purified by size-exclusion chromatography, concentrated, and flash-frozen. **b**, SDS-PAGE analysis of input cytoplasmic human Pol II. **c**, SDS-PAGE analysis of recombinant human assembly/transport factors for Pol II spike-in. **d**, SDS-PAGE analysis of cytoplasmic human Pol II spiked with assembly/transport factors and purified by size-exclusion chromatography. **e**, SDS-PAGE analysis of BS3-crosslinked cytoplasmic human Pol II spiked with assembly/transport factors. Inp: Concentrated spiked cytoplasmic human Pol II. BS3: BS3 crosslinked spiked cytoplasmic human Pol II. Gels when labelled: Molecular weight markers (left); Pol II subunits and recombinant assembly/transport factors labelled (right).

**Extended Data Fig. 2:**
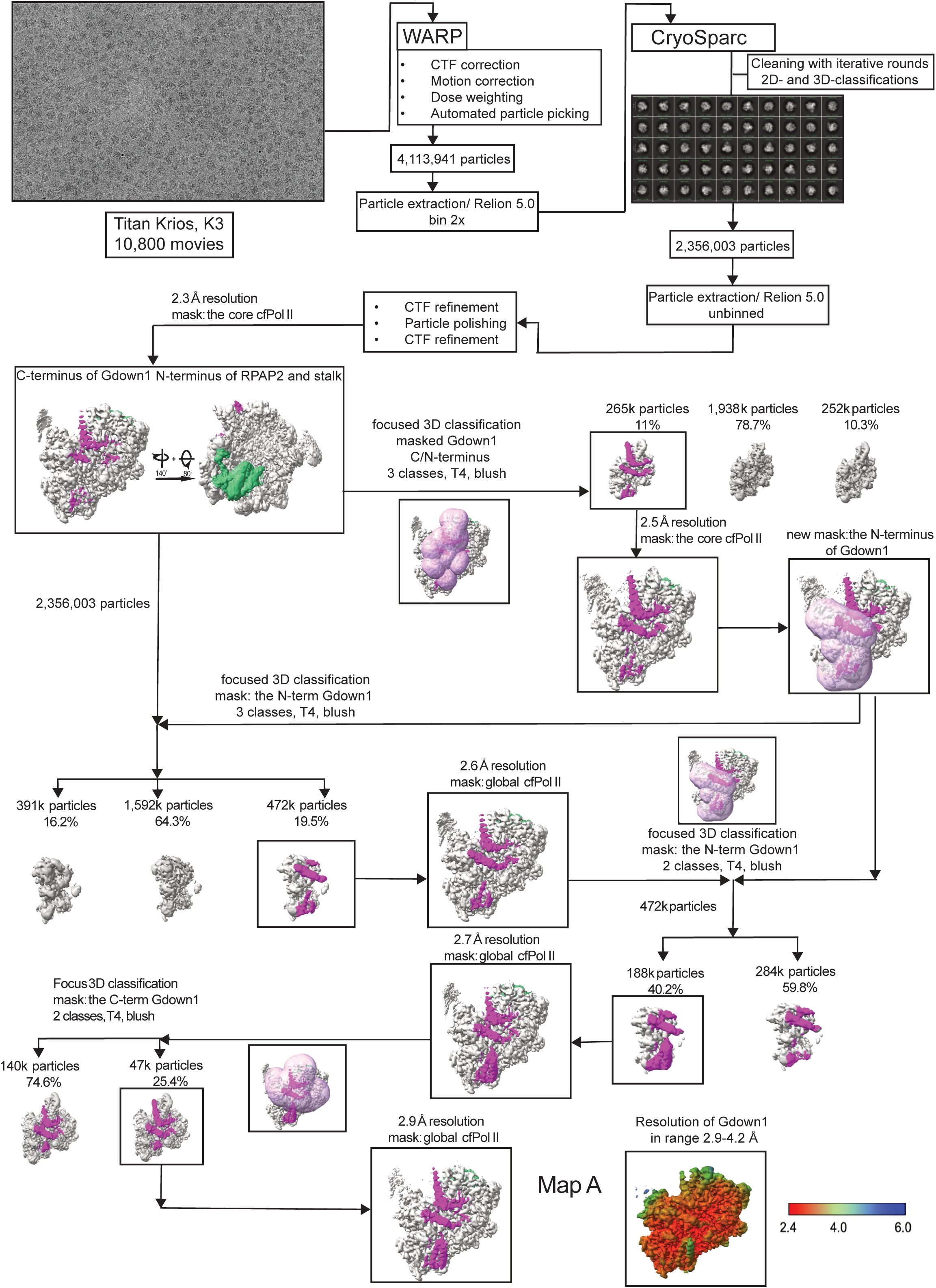
Cryo-EM analysis of cytoplasmic human Pol II complexed with assembly factors RPAP2, GPN1, GPN3, and Gdown1 (Map A; Gdown1-focused) The cryo-EM processing workflow begins with a representative micrograph visualized in Warp^39^. The processing tree outlines 3D classification steps and focused refinements that improved density for Gdown1 domains. Particle classes exhibiting optimal Gdown1 density (highlighted in pink and boxed) were selected for 3D refinement. Resulting refined maps are displayed adjacent to their corresponding 3D classification classes. Distinct extra densities are highlighted as follows: Gdown1 at RPB2 protrusion (magenta) and RPAP2 at RPB5 (green). Mask regions for Gdown1-focused 3D classification appear as rosy-brown transparent surfaces. Local resolution of Gdown1 was calculated in RELION 5.0^40,41^ and visualized in ChimeraX^44^.

**Extended Data Fig. 3:**
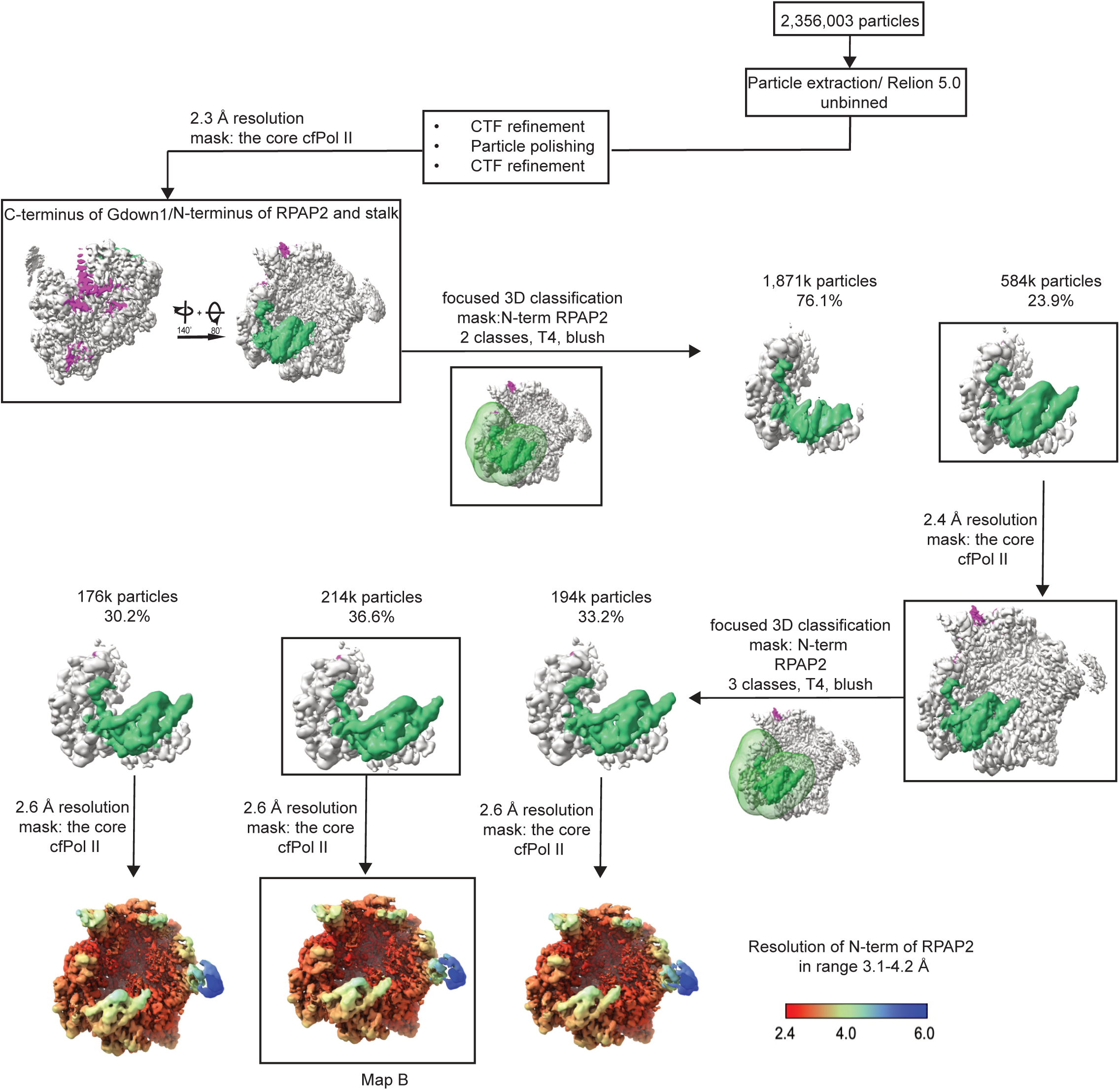
Cryo-EM analysis of cytoplasmic human Pol II complexed with assembly factors RPAP2, GPN1, GPN3, and Gdown1 (Map B; RPAP2-focused) The cryo-EM processing workflow outlines 3D classification steps and focused refinements that improved density for the N-termini of RPAP2. Particle classes exhibiting optimal RPAP2 density (highlighted in green and boxed) were selected for 3D refinement. Resulting refined maps are displayed adjacent to their corresponding 3D classification classes. Local resolution maps for RPAP2 were calculated in RELION 5.0^40,41^ and visualized in ChimeraX^44^.

**Extended Data Fig. 4:**
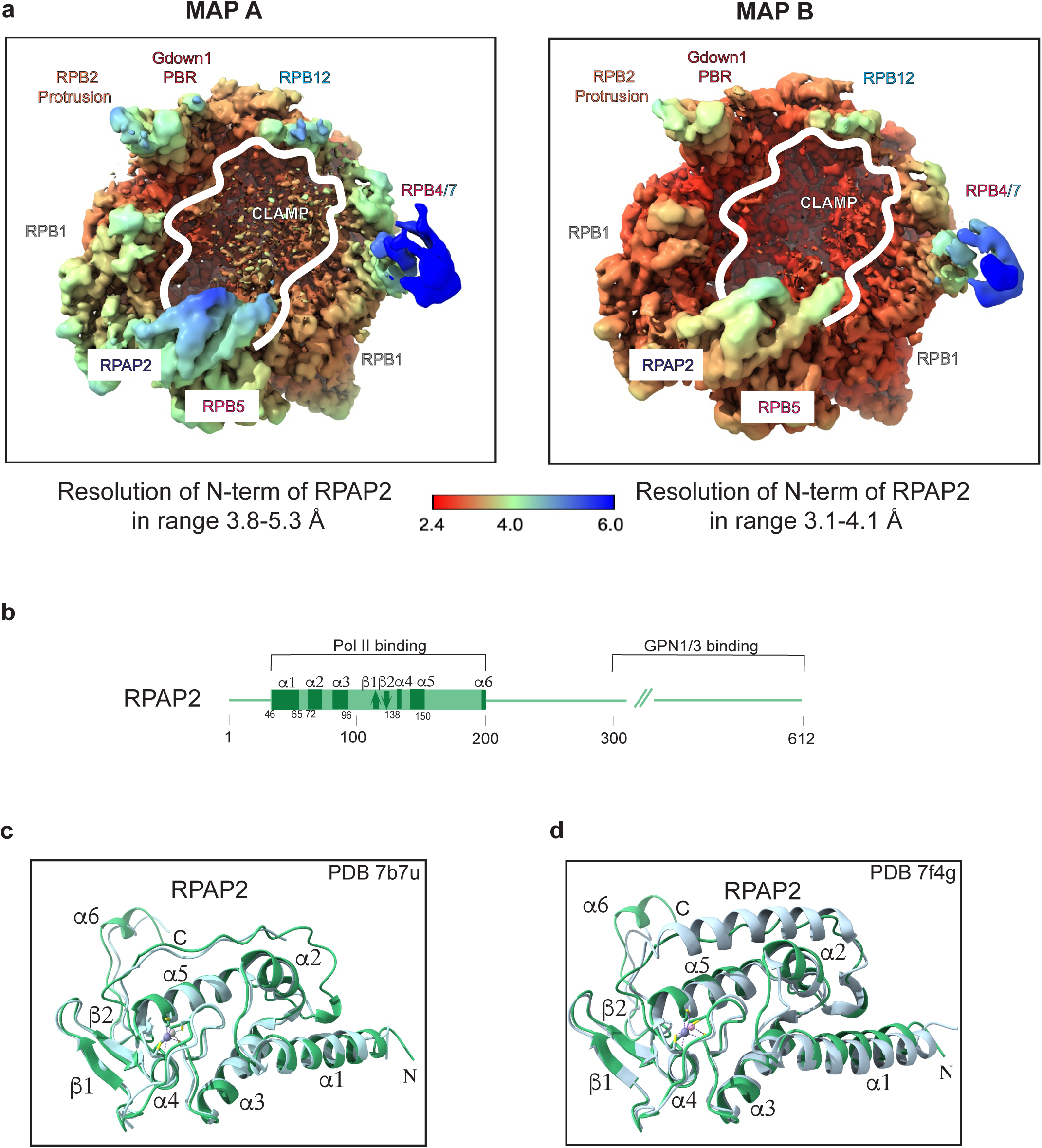
Structure details of cfPol II. **a**, Local resolution comparison of the N-terminal RPAP2 region (residues 46–197) between Map A (Gdown1-focused) and Map B (RPAP2-focused). Local resolution maps were calculated in RELION 5.0^40,41^ and visualized in ChimeraX^44^. **b**, Primary architecture of human RPAP2. The confidently modelled N-terminal region (residues 46-197) is shown in light green boxes with secondary structure elements (α-helices: rectangles; β-sheets: arrows) indicated in dark green. The disordered C-terminal region (residues 198–612) is represented by a green line and could not be modelled in our cryo-EM maps. **c**, Structural alignment of our RPAP2 ribbon model (this study) with published RPAP2 structures: Left - PDBs 7B7U^27^; Right - PDB 7F4G^28^.

**Extended Data Fig. 5.**
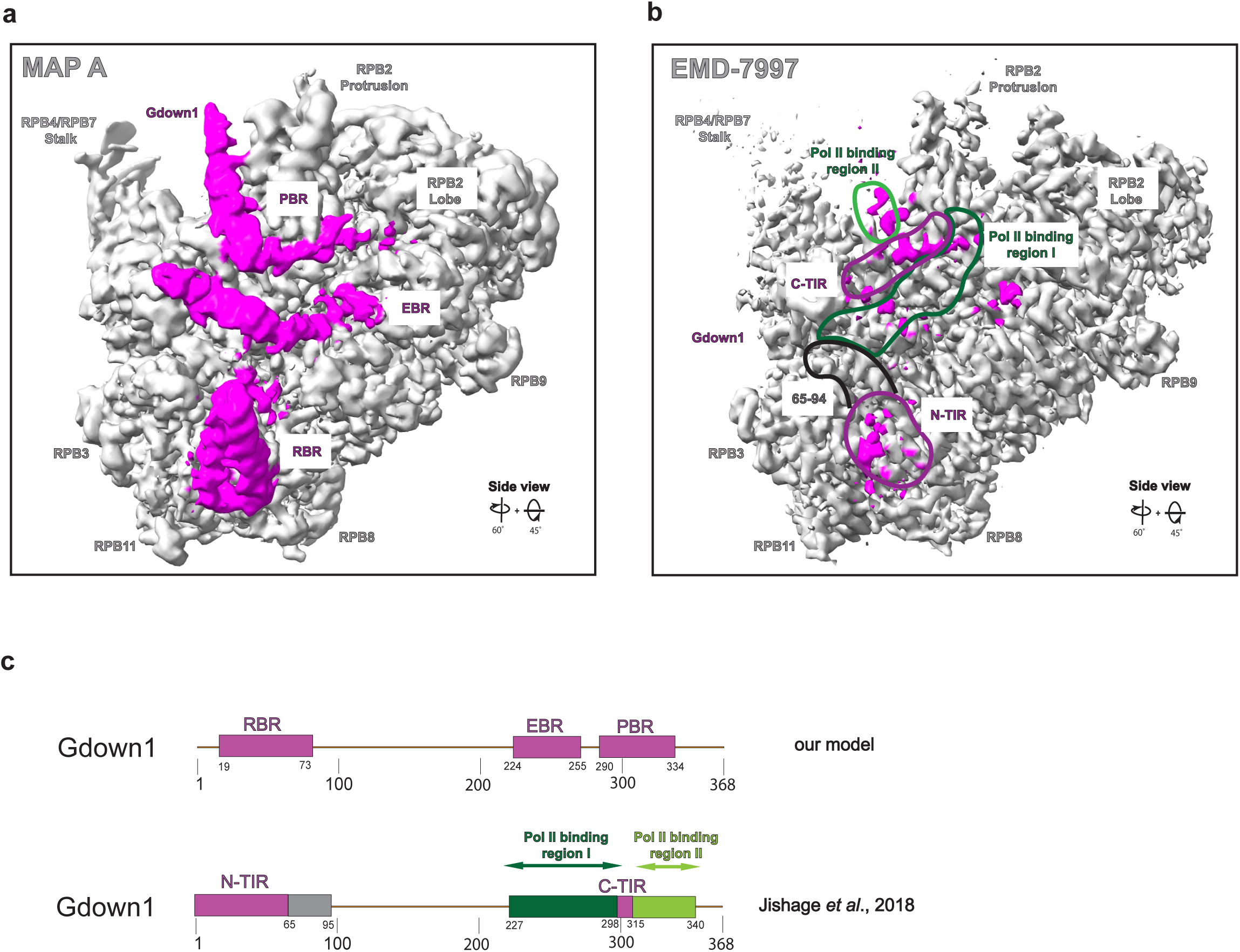
Cryo-EM density comparison of Gdown1 in Map A versus EMDB-3797. **(a)** *Map A:* Gdown1 cryo-EM density (magenta) with fitted atomic model from Map A. Regions binding RPB2, RPB10, and RPB3 are labeled according to Fig. 2a,b designations (RBR, EBR, PBR). **(b)** *EMDB-3797* (Jishage *et al*., 2018^30^): Gdown1 density (magenta) fitted with the same atomic model as in (a). Binding regions labeled per Jishage *et al*., 2018^30^ (Fig. 4h). **(c)** Domain architecture comparison of human Gdown1 between our model and Jishage *et al*., 2018^30^.

**Extended Data Fig. 6.**
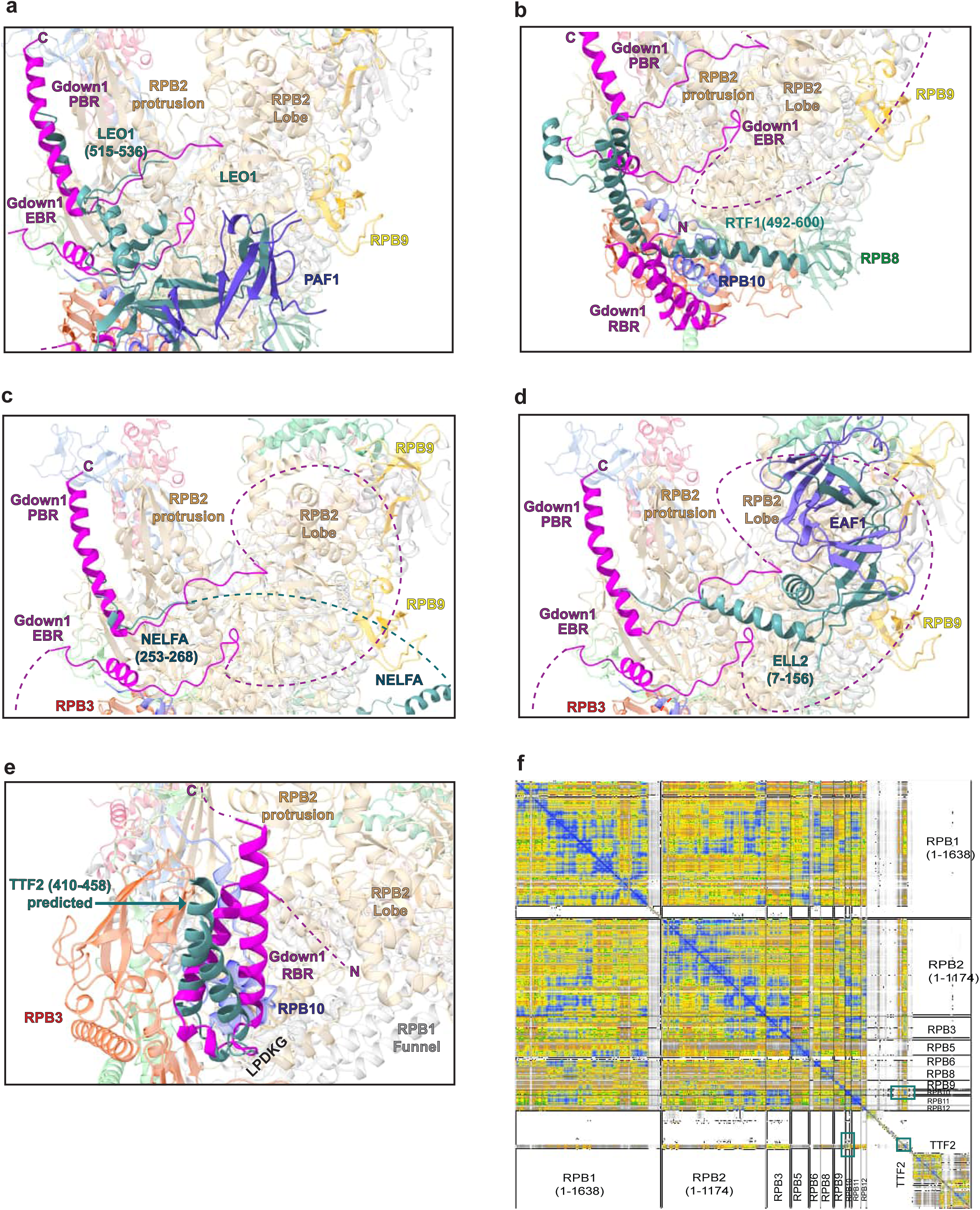
Steric clashes between Gdown1 and transcription elongation and pausing factors, as well as between Gdown1 and the transcription termination factor TTF2. **(a)** Gdown1 PBR (magenta) sterically clashes with LEO1 (teal; PDB: 9RTT^53^) **(b)** Gdown1-RPB2 interface conflicts with RTF1 (teal; PDB: 6TED^54^), an elongation/pausing/chromatin regulator. **(c)** Gdown1 obstructs NELFA tentacle domain binding (teal; PDB: 8UI0^55^). **(d)** Gdown1 PBR/EBR regions block ELL2-EAF1 complex binding (ELL2: teal; EAF1: blue; PDB: 7OKX^56^). **(e)** Superposition of TTF2-Pol II AlphaFold model reveals clashes between Gdown1-RBR (RPB10 interface) and TTF2 residues 410–458. Pol II shown at 80% transparency. **(f)** *Left:* Predicted Aligned Error (PAE) plot for TTF2-Pol II AlphaFold model. Red boxes indicate TTF2 new fold (residues 410–458) binding RPB10. *Right:* Cartoon representation colored by pLDDT confidence. *Note:* Higher PAE values indicate lower structural confidence.

