## Extended Data Table 1 for "Structure of human cytoplasmic Pol II complex explains global transcription repression by Gdown1"

**Extended Data Table1. Mass spectrometry analysis of endogenous cfPol II complex**

| Protein complex | Total spectra | Sequence coverage (%) | Mw (kDa) |
|---|---|---|---|
| human Pol II | | | |
| N-term 6xHisTwinStrep_RPB1 | 9002 | 77 | 223 |
| RPB2 | 6734 | 91 | 134 |
| RPB7 | 1370 | 100 | 19 |
| RPB3 | 1218 | 94 | 31 |
| RPB4 | 1118 | 80 | 16 |
| RPB9 | 973 | 87 | 15 |
| RPB5 | 928 | 85 | 25 |
| RPB8 | 534 | 91 | 17 |
| RPB10 | 517 | 94 | 8 |
| RPB11 | 428 | 100 | 13 |
| RPB6 | 249 | 55 | 14.5 |
| RPB12 | 142 | 62 | 7 |
| Assembly and transport factors | | | |
| RPAP2 | 1490 | 80 | 70 |
| Gdown1 | 1334 | 92 | 42 |
| GPN3 | 580 | 68 | 33 |
| RECQL5 | 537 | 76 | 109 |
| GPN1 | 496 | 81 | 42 |
| RUVB2 | 224 | 79 | 51 |
| RUVB1 | 214 | 82 | 50 |
| HS71A | 152 | 66 | 70 |
| HSPA8 | 127 | 56 | 71 |
| SLC7A6OS (hIWR1) | 58 | 45 | 35 |
| URI1 | 58 | 44 | 60 |
| RPAP3 | 36 | 41 | 75.7 |
| HSP90 | 21 | 14 | 84.6 |
| Cytoplasmic contaminants | | | |
| Acetyl_CoA carboxylase 1 | 6394 | 95 | 266 |
| Acetyl_CoA carboxylase 2 | 898 | 63 | 277 |
| Tubulin cluster Beta | 1286 | 85 | 50 |
| Tubulin cluster alpha | 672 | 97 | 50 |
| UBR4 | 307 | 33 | 574 |
| KRT10 | 113 | 53 | 59 |
| KRT2 | 101 | 61 | 65 |
| RPS3 (ribosomal protein) | 80 | 81 | 27 |
| RPS4X (ribosomal protein) | 61 | 52 | 30 |
| Mitochondrial contaminants | | | |

| | | | |
|---|---|---|---|
| PCCA | 247 | 67 | 80 |
| MCCB | 198 | 76 | 61 |
| MCCA | 172 | 62 | 80.5 |
| PCCB | 194 | 81 | 58 |
| Nuclear proteins/complexes | | | |
| Integrator: | | | |
| INT1 | 407 | 54 | 244 |
| INT4 | 203 | 62 | 108 |
| INT7 | 183 | 63 | 107 |
| INT2 | 140 | 49 | 134 |
| INT3 | 136 | 49 | 118 |
| INT10 | 135 | 57 | 82 |
| INTS13 | 129 | 57 | 80 |
| INT5 | 113 | 52 | 108 |
| INT6 | 105 | 46 | 100 |
| INT8 | 86 | 41 | 113 |
| INT9 | 84 | 54 | 73.8 |
| INT14 | 60 | 61 | 57 |
| SPT6 | 97 | 36 | 199 |
| Mediator | | | |
| Med23 | 64 | 20 | 156.5 |
| DNA damage | | | |
| WRNIP1 | 137 | 58 | 72 |
| FBX21 | 146 | 51 | 70.3 |
| SKP1 | 68 | 89 | 19 |

**Table 1.**
Proteins were identified by LC-MS/MS from BS3 cross-linked sample of endogenous cfPol II. Total spectra, sequence coverage (%), and molecular weight (Mw) are shown. Proteins are grouped by functional class.
