## Extended Data Table 2 for "Structure of human cytoplasmic Pol II complex explains global transcription repression by Gdown1"

### Extended Data Table2. Cryo-EM data acquisition, processing and refinement statistics

| **Data collection and processing** | **CMPol II-GDown1-RPAP2 MAP A** |
|---|---|
| Magnification | 81,000x |
| Voltage (kV) | 300 |
| Electron exposure (e–/Å2) | 39.91 |
| Defocus range (μm) | 0.6-2 |
| Pixel size (Å) | 1.05 |
| Initial particle images (no.) | 11,000 |
| PDB code | PDB: XXXX |
| Map code | EMDB: EMD-XXXXX |
| Symmetry imposed | C1 |
| Map resolution (Å) at FSC = 0.143 | 2.9 |
| Map resolution range (Å) | 2.9-5.0 |
| **Refinement** | |
| Initial models used (PDB code) | 7ozn |
| **Model composition** | |
| Non-hydrogen atoms | 29,491 |
| Protein residues | 3,668 |
| Ligands | Zn: 6 |
| **Mean B factors ($Å^2$)** | |
| Protein | 90.77 |
| Ligand | 155.53 |
| **R.m.s. deviations** | |
| Bond lengths (Å) | 0.004 |
| Bond angles (°) | 0.830 |
| **Validation** | |
| MolProbity score | 1.67 |
| Clashscore | 6.35 |
| Poor rotamers (%) | 0 |
| **Ramachandran plot** | |
| Disallowed (%) | 0.00 |
| Allowed (%) | 4.56 |
| Favored (%) | 95.44 |
